# High-Dimensional Imaging of Vestibular Schwannoma Reveals Distinctive Immunological Networks Across Histomorphic Niches in *NF2*-related Schwannomatosis

**DOI:** 10.1101/2024.04.09.588593

**Authors:** Adam P. Jones, Michael J. Haley, Grace E. Gregory, Cathal J. Hannan, Ana K. Simmons, Leoma D. Bere, Daniel G. Lewis, Pedro Oliveira, Miriam J. Smith, Andrew T. King, D Gareth R. Evans, Pawel Paszek, David Brough, Omar N. Pathmanaban, Kevin N. Couper

## Abstract

*NF2*-related Schwannomatosis (*NF2* SWN) is a rare tumour-predisposition syndrome characterised by the growth of multiple central and peripheral nervous system neoplasms. The drivers of *NF2* SWN are pathogenic variants in the tumour suppressor gene *NF2*, encoding the protein Merlin, leading to development of bilateral vestibular schwannoma (VS) in >95% of patients. VS tumours are characterised by infiltration of myeloid cells and lymphocytes, highlighting the potential of immunotherapy for VS. However, the immunological landscape in VS and the spatial determinants within the tumour microenvironment that shape the trajectory of disease are presently unknown. In this study, to elucidate the complex immunological networks across VS, we performed imaging mass cytometry (IMC) on clinically annotated VS samples from *NF2* SWN patients. We reveal the heterogeneity in neoplastic cell, myeloid cell and T cell populations that co-exist within VS, determining that the cellular composition of VS tumours is independent of *NF2*-SWN genetic severity. We show that distinct myeloid cell and Schwann cell populations exist within varied spatial contextures across characteristic Antoni A and B histomorphic niches. Interestingly, we show that T-cell populations associate with tumour-associated macrophages (TAMs) in Antoni A regions, seemingly limiting their ability to interact with tumorigenic Schwann cells. This spatial landscape is altered in Antoni B regions, where T-cell populations appear to interact with PD-L1+ Schwann cells. We also demonstrate that prior bevacizumab treatment (VEGF-A antagonist) preferentially reduces alternatively-activated TAMs, whilst enhancing CD44 expression, in bevacizumab-treated tumours. Together, we describe niche-dependent modes of T-cell regulation in *NF2* SWN VS, indicating the potential for microenvironment-altering therapies for VS.

**Teaser:** Imaging mass cytometry and spatial omic analyses illustrate spatially-distinct regions of T-cell regulation in vestibular schwannoma.

## Introduction

Bilateral vestibular schwannomas (VS) are the pathognomonic hallmark of the tumour predisposition syndrome *NF2*-related Schwannomatosis (*NF2* SWN)^1^. *NF2* SWN is caused by germline or tissue mosaic pathogenic variants (PV) in the tumour suppressor gene *NF2*, encoding a Moesin-Ezrin-Radixin-like protein (termed Merlin), which results in development of bilateral VS in >95% of afflicted patients^2–4^. The severity phenotype of *NF2* SWN^5^ depends, in part, on the type of PV in the *NF2* gene, with whole gene deletions and missense variants resulting in a milder phenotype (later onset of symptoms and lower tumour burden), compared with truncating and frameshift variants, which cause the severe phenotype characterised by early onset of symptoms and high tumour burden^6–9^.

Whilst VS are benign intracranial tumours, their development and treatment (surgical and radiation^10,11^) often cause severe morbidity such as sensorineural hearing loss and facial weakness, and left unchecked, may lead to brainstem compression and hydrocephalus, requiring urgent neurosurgical intervention^12–14^. Currently, there are no approved drugs for the treatment of *NF2* SWN or VS. Multiple studies have trialled the use of bevacizumab, an inhibitor of angiogenesis, in *NF2* SWN-related VS and found a reduction in tumour growth and hearing stabilisation or improvement^15–18^, but treatment toxicity such as hypertension, proteinuria and renal impairment are common, and many individuals treated with bevacizumab become refractory, resulting in uncontrolled growth and progressive hearing loss^19–21^. The current lack of alternative treatments for *NF2* SWN-related VS is a consequence of our limited understanding of the biology of *NF2*-SWN-related VS tumours.

Whilst the tumour architecture of VS has been defined at the pathological level with the discovery of hypercellular Antoni A regions, and less cellular Antoni B regions^22^, the cellular landscape in Antoni A and B regions, in particular the compartmentalisation and activities of immune cells within these distinct regions, has yet to be investigated. Notably, VS tumours harbour significant numbers of macrophages and T-cells^23,24^. It has been suggested that macrophages in VS have a suppressive tumour-associated macrophage (TAM) phenotype^25^ and the abundance of TAMs has been linked with VS tumour growth rate, potentially through vascular endothelial growth factor (VEGF) expression and promotion of angiogenesis^25–27^. A recent transcriptomic study identified signatures associated with immune enrichment and CD8^+^ T-cell senescence in rapidly progressing VS^28^. Despite these observations, there is a paucity of data describing how macrophages, T cells and neoplastic Schwann cells compartmentalise and interact within the VS tumour microenvironment (TME) and histomorphic niches.

In this study we have employed Hyperion imaging mass cytometry (IMC) to spatially map the immunological landscape in Antoni A and B regions of *NF2* SWN VS tumours. We found that the spatial interactome of Antoni A and B regions differ, whereby the perivascular niche in Antoni B regions is disrupted and loses connectivity to supportive immune cells, suggesting potential vascular degeneration in these regions. T-cells also have distinct spatial profiles in Antoni A and B regions, highlighting two niche-dependent regulatory networks in which T-cells exist within VS. Notably, we identified that disease severity and *NF2* pathogenic variants did not appear to have any significant influence on the TME, whilst Bevacizumab-treatment was associated with a significant increase in CD44+ Schwann cells, suggesting a potential increase in matrix remodelling within these tumours. Collectively, our results provide new insight into the immunological configuration of *NF2* SWN VS tumours and suggest tumour region-specific pathways that are potential therapeutic targets for the disease.

## Methods

### Tissue samples and ethics

16 retrospective *NF2* SWN VS cases (see Table 1 for clinical information), in formalin-fixed paraffin-embedded (FFPE) blocks, were accessed through the cellular pathology department at the Manchester Centre for Clinical Neurosciences, Salford Royal Hospital (SRH) and approved by the Health Research Authority (HRA) and Health and Care Research Wales (HCRW) for research (REC: 20/NW/0015, IRAS ID: 274046). Haematoxylin and eosin (H&E) stained slides were generated by pathology at SRH for each case and regions of interest (ROI) were selected by a neuropathologist based on identification of typical VS pathological features (Antoni A regions, Antoni B regions, immune infiltration, abnormal vasculature). Once identified, 2mm^2^ cores of each ROI were removed from each block and re-formatted into tissue microarrays (TMA).

**Table 1:** Case info.

| Case | Sex | Age at radiological diagnosis | Age at surgery | NF2 pathogenic variant (NM_000268.4) | Disease severity | Prior treatments | Bevacizumab treatment |  |  |  |
| --- | --- | --- | --- | --- | --- | --- | --- | --- | --- | --- |
|  |  |  |  |  |  |  | Dose | Regime | Duration | Cessation prior to surgery |
| NF2VS1 | M | 20 | 27 | Unknown (likely mosaic) | Mild | None |  |  |  |  |
| NF2VS2 | M | 57 | 59 | c.999+1G>T, p.(?) | Mild | None |  |  |  |  |
| NF2VS3 | F | 47 | 52 | Whole gene deletion | Mild | Resection |  |  |  |  |
| NF2VS4 | F | 29 | 35 | c.1604T>C, p.(Leu535Pro) | Mild | None |  |  |  |  |
| NF2VS5 | F | 44 | 50 | c.1229_1241del, p.(Gln410Profs*12) | Severe | None |  |  |  |  |
| NF2VS6 | M | 45 | 53 | c.516+1G>T, p.(Leu361Cysfs*3) | Moderate | None |  |  |  |  |
| NF2VS7 | M | 22 | 22 | Deletion of promotor to exon 1 | Mild | None |  |  |  |  |
| NF2VS8 | F | 26 | 26 | c.1080del, p.(Leu361Cysfs*3) | Severe | None |  |  |  |  |
| NF2VS9 | M | 12 | 17 | c.1396C>T, p.(Arg466*) | Severe | Bevacizumab | 10mg/kg | Every 2 weeks | 5.0 years | 8 weeks |
| NF2VS10 | M | 15 | 20 | Deletion of exons 7 to 14 | Moderate | Bevacizumab | 10mg/kg | Every 2 weeks | 3.8 years | 12.7 weeks |
| NF2VS11 | M | 31 | 31 | c.265G>T, p.(Glu89*) | Severe | None |  |  |  |  |
| NF2VS12 | M | 29 | 29 | c.1606C>T, p.(Gln536*) | Moderate | None |  |  |  |  |
| NF2VS13 | F | 32 | 33 | Deletion of exon 15 | Moderate | None |  |  |  |  |
| NF2VS14 | F | 32 | 34 | c.1021C>T, p.(Arg341*) | Severe | None |  |  |  |  |
| NF2VS15 | F | 22 | 29 | Deletion of intron 1 to exon 17 | Moderate | Resection |  |  |  |  |
| NF2VS16 | F | 24 | 33 | c.169C>T, p.(Arg57*) | Severe | Bevacizumab | 7.5mg/kg | Every 3 weeks | 4.4 years | 13.7 weeks |

### H&E staining and imaging

Case blocks/TMAs were sectioned at 5μm thickness using a microtome. The 5μm tissue sections were deparaffinised in xylene for 5 mins and rehydrated through an alcohol gradient (100-70% ethanol, 1 min each, then into water). Hydrated sections were incubated in haematoxylin for 3 mins, followed by washing under running water for 2 mins. Sections were then dipped in 1% hydrochloric acid-ethanol solution for 10 secs, followed by washing under running water for 3 mins. Sections were then blued using Scott’s tap water for 20 secs, followed by washing under running water for 5 mins. Next, sections were submerged in 70% ethanol for 10 secs, before being counterstained with Eosin Y for 30 secs. Sections were dehydrated through an alcohol gradient (70-100% ethanol, 30 secs each), then xylene for 3 mins. Slides were covered slipped with DPX mountant. H&E-stained slides were imaged on the Olympus VS200 Slide Scanner (Olympus LifeScience) and digitally visualised using CaseViewer (v2.4, 3DHISTECH).

### IMC antibody conjugation, optimisation and validation

The IMC antibody panel used for this study is outlined in Table 2: sources of all antibodies are outlined in Supplementary Table 1. Antibodies were initially tested by immunofluorescence (IF) on several positive control tissues (such as secondary lymphoid tissues and tumour samples). Antibodies were conjugated to metal isotopes using the MaxPar Multimetal Labelling Kit (Standard BioTools), whereby metal isotopes were loaded onto chelating polymers, and subsequently bound to the antibody. Antibodies conjugated to platinum isotopes underwent a different conjugation protocol^29^.

**Table 2:**
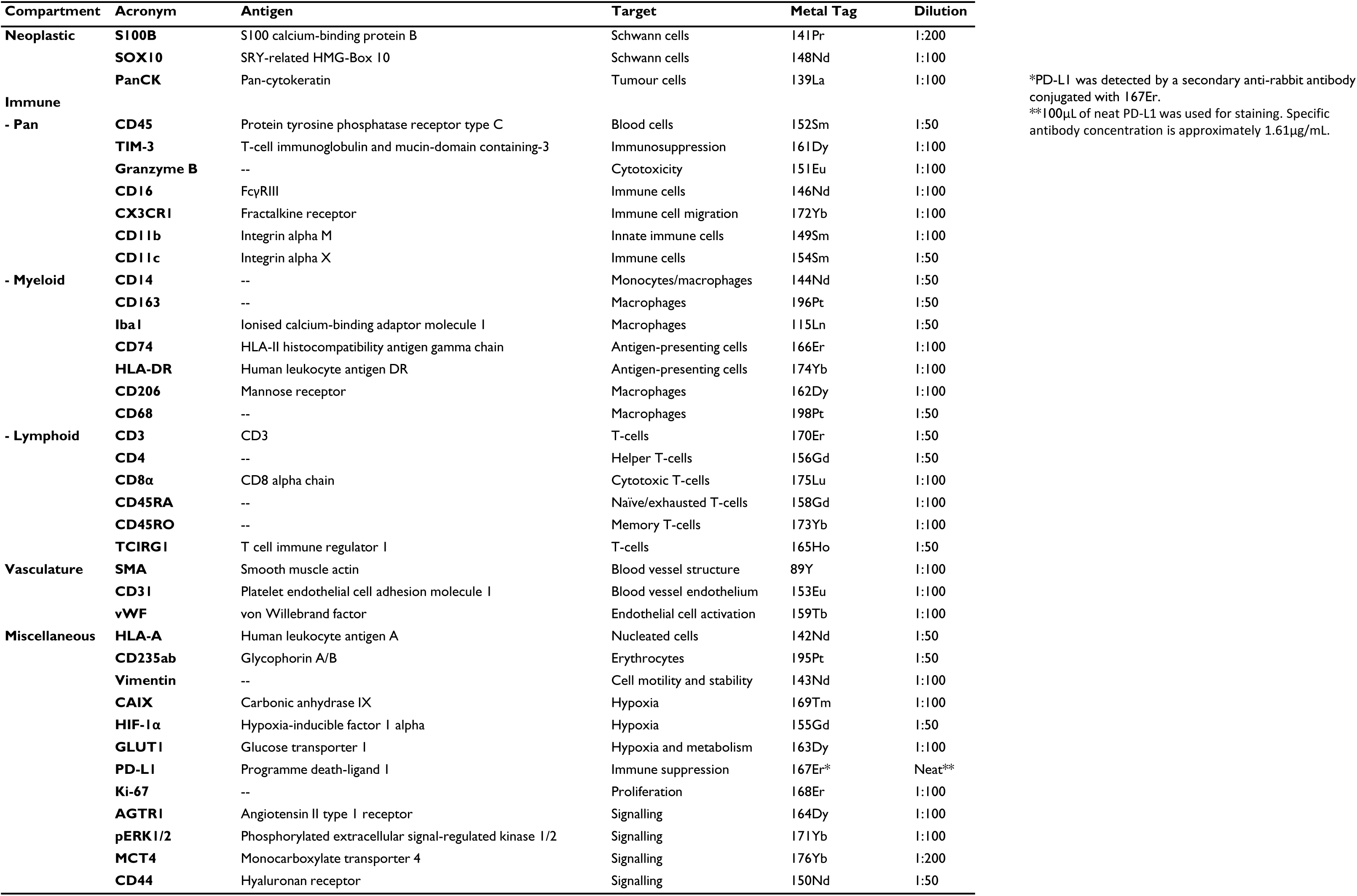
IMC panel.

### IMC staining

Staining for IMC was performed according to Standard BioTools optimised protocols for FFPE tissues^30^. TMA blocks were sectioned at 5μm thickness with a microtome, deparaffinised in fresh xylene for 10 mins, and rehydrated through an ethanol gradient (100– 50%, 1 min per grade), and placed in distilled water. Next, tissue sections underwent antigen retrieval by incubation at 96°C in Tris-EDTA (pH 8.5) for 30 mins. Once cooled to 70°C, tissue sections were washed in PBS (0.05% Tween) and blocked with 3% bovine serum albumin (BSA) for 45 mins at room temperature (RT). Next, sections were incubated with diagnostic-grade anti-PD-L1 for 1 hour at RT, washed, and a secondary anti-rabbit IgG-167Er antibody was added for 1 hour at RT. Slides were then washed and the incubated with the remaining IMC antibodies (dilutions outlined in Table 2, diluted in PBS and 0.5% BSA) and incubated overnight at 4°C. The following day, slides were washed with 0.1% Triton X-100 PBS, and incubated with Cell-ID Intercalator-Iridium (1:400 in PBS, Standard BioTools), for 30 mins at RT, washed with deionised water (Merck Millipore), and dried overnight.

### IMC image acquisition

Raw IMC images were acquired using a Hyperion imaging mass cytometer coupled with a Helios time-of-flight mass cytometer (CyTOF, Standard BioTools). The previously neuropathologist-defined ROIs (Antoni A and B regions) were selected at 1000×1000μm, and the selected tissue regions were laser ablated over several consecutive days, in a rastered pattern at 1μm^2^ pixel resolution, and at a frequency of 200Hz. The resultant plume following tissue ablation was passed through a plasma source, ionising it into constituent atoms. CyTOF then differentiated the signal from each of the metal-conjugated antibodies, and images of each antibody were reconstructed based on abundance of each metal at each pixel. Images for each antibody were then exported from the raw data as TIFFs using MCD Viewer (Standard BioTools). Acquired ROIs were screened for abnormalities during image generation, and were excluded accordingly.

### Denoise clean-up and single-cell segmentation of IMC images

Raw IMC images were subjected to a denoise pipeline according to authors instructions^31^. Following denoising, single-cell data was mined from IMC images using an established protocol^32^. Stacks of TIFF images were exported from MCD files of each acquired ROI, with each channel corresponding to a metal isotope-conjugated antibody. Ilastik^33^ was then applied to produce pixel probability maps that distinguished between nuclear, cytoplasmic and background pixels. These maps were then converted into cell segmentation masks, where individual cell boundaries were defined, and masks were applied to each antibody channel, producing single-cell expression data for each channel, including the spatial location of each cell within the acquired ROIs.

### Single-cell analyses of IMC data

The single-cell expression data was interrogated in Python using packages devised to analyse single-cell data (Scanpy vX^34^) and spatial data (Squidpy vX^35^ and ATHENA vX^36^). Mean cell intensity of each marker was normalised to the 99.9^th^ percentile of its expression, and batch corrected by case using batch balanced k nearest neighbours (BBKNN) alignment^37^. Leiden clustering^38^ was used to identify cell populations. Initial Leiden populations were divided into 4 main groups (Schwann cells, myeloid cells, lymphoid cells and vasculature) based on canonical markers of each group. These broad classified cell clusters were reclustered at higher resolution to identify more granular cell sub-populations (supplementary Fig 1B), an approach performed in other high dimensional imaging studies^39^. These population were then re-defined on the original Leiden and annotated based upon marker expression of known cell types and states. Single-cell metrics for each population were extracted for each ROI, and used to calculate cell abundance.

### Spatial omics analyses

Spatial analyses were performed according to the Spatial Omics Oxford (SpOOx) analysis pipeline^40,41^. Significant associations between different cell types were assessed by cross pair correlation functions (cross-PCF), which was used to measure direct cell-cell pairwise partners within 20µm of each cell. Adjacency cell network (ACN) analyses were generated by computing pairwise statistics between combinations of cell types, ‘A’ and ‘B’. This was done by calculating the amount of ‘A’ cells in contact with ‘B’ cells, and then subsequently calculating the proportion of ‘B’ cells in contact with ‘A’ cells within each ROI, for each cell.

### Cellular neighbourhood analyses

Derivation of cellular neighbourhoods was performed using CellCharter^42^. Initially, cells were sorted into neighbourhood aggregates, and then clustered by Gaussian mixture modelling (GMM). To determine the optimal number of clusters, *K*, GMM clustering was run multiple times (10 for this study) and the highest stability clustering *n* was selected in accordance with the Fowlkes–Mallows Index (FMI). All cells were then sorted into spatial clusters. Next, proximity analysis was performed to assess the relative arrangement of spatial clusters within ROIs, indicating neighbourhood enrichment between clusters.

### Statistics

Statistical tests were performed using GraphPad Prism (v9.5.1). Data distribution was assessed by Shapiro-Wilk normality test. Differences in case-averaged cell abundances between two groups (Antoni A versus Antoni B; naïve versus bevacizumab) were performed employing case averages using unpaired t-test or Mann Whitney U test. For differences between three groups (disease severity), cell abundances were compared employing case averages using a one-way ANOVA with Tukey’s multiple comparisons test or Kruskal-Wallis test with Dunnett’s T3 multiple comparisons test.

## Results

### Imaging mass cytometry highlights the intra and inter-heterogeneity across histomorphic niches in *NF2* SWN-related vestibular schwannoma

To explore the single-cell and spatial heterogeneity of VS in *NF2* SWN, 13 treatment naïve VS cases from individuals with *NF2* SWN (Table 1) were analysed within a programme of work outlined in Figure 1A. H&E tissue sections were initially evaluated by a neuropathologist, annotated as Antoni A or B regions (Figure 1B), and incorporated into a 73-core TMA, with 2-8 cores per case. Sections were stained with a panel of 40 metal-conjugated antibodies (Table 2), which was designed to allow interrogation of Schwann cell, myeloid cell, lymphoid cell, and vascular-related populations. Representative images showing staining of Schwann cells (S100B), macrophages (Iba1), T-cells (CD8), stroma (SMA) and proliferative cells (Ki-67) in Antoni A and B regions are shown in Figure 1B. Following single-cell segmentation using Ilastik, we analysed the expression of canonical markers on cells by UMAP. This showed that markers specific to Schwann cell (S100B+, SOX-10+), myeloid (Iba1+, CD68+, CD163+), lymphoid (CD8+, CD4+), and vascular-related (SMA+, CD31+) populations were segregated into distinct clusters (supplementary Fig 1A). Using Leiden clustering (outlined in supplementary Fig 1B) we identified 23 distinct populations across the 13 VS cases (Fig 1C and supplementary Fig 1C). Of note, 7 Schwann cell populations, 7 myeloid populations, 4 lymphoid (T-cell) populations, and 2 vascular populations were identified, with the remainder classified as erythrocytes, proliferating cells, and other cells that could not be clearly resolved with the panel (other) (Fig 1C); interestingly, these data suggest that myeloid cells and Schwann cells exist on a spectrum of activation and polarization states in VS, indicating both pro-inflammatory and regulatory phenotypes.

**Figure 1:**
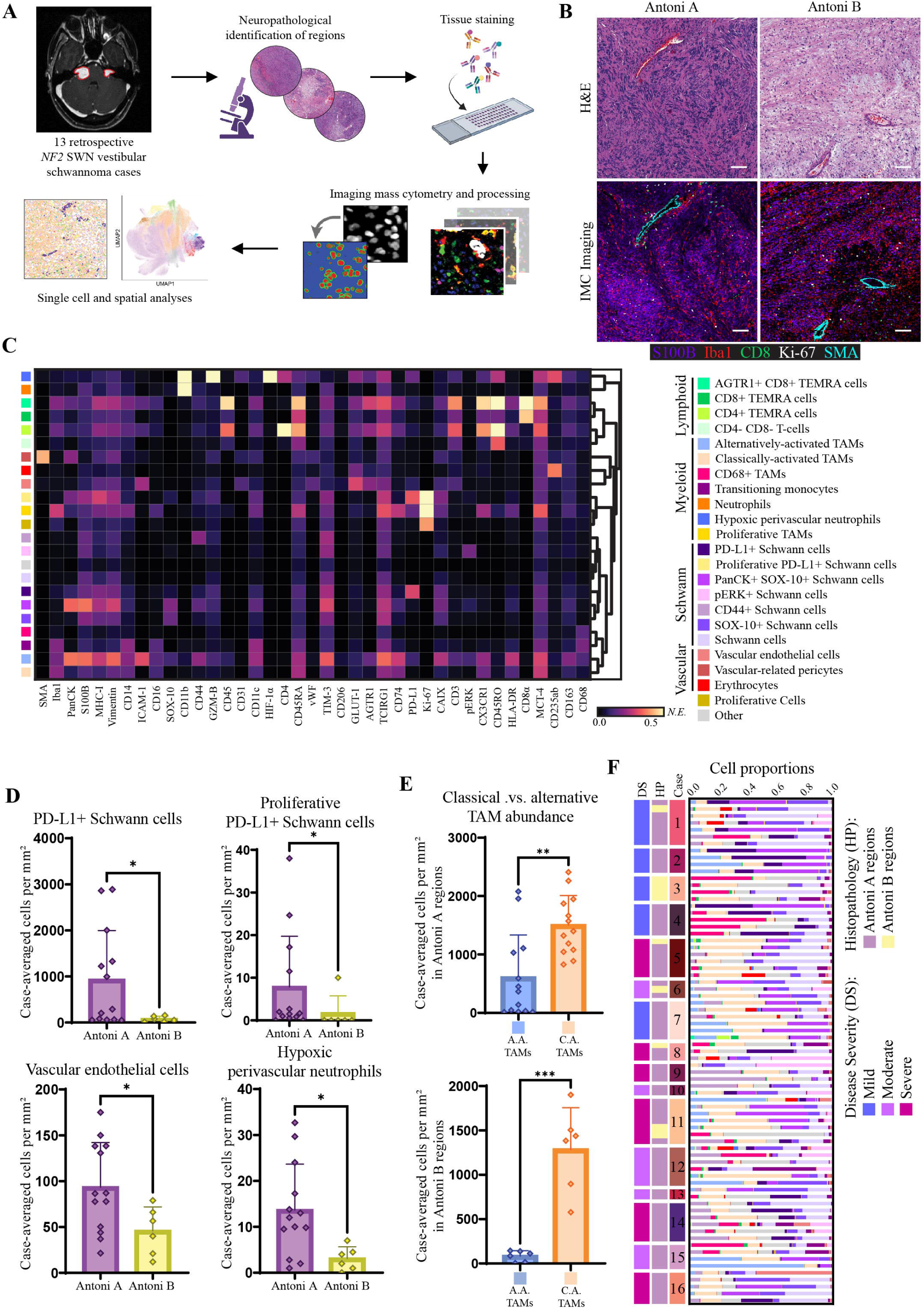
Imaging mass cytometry (IMC) workflow and generation of the single-cell atlas in *NF2* SWN-related vestibular schwannoma. **A:** Schematic of IMC workflow for investigating *NF2* SWN-related VS: niche identification, tissue processing and staining, data acquisition and processing of raw data, single-cell segmentation, and downstream analyses. **B:** From left to right, sequential haematoxylin and eosin (H&E) and IMC images of Antoni A and Antoni B region, visualising the core Schwann cell (S100B; purple), macrophage (Iba1, red), T-cell (CD8α; green), vascular (SMA; cyan), and proliferation (Ki-67; white) markers. Scale bar representative of 100μm. **C:** Heatmap detailing expression pattern of all single-cell populations identified in IMC analysis via Leiden clustering. Scale bar indicating normalised expression (N.E.). **D:** Comparison of case-averaged abundance of significantly different cell populations across Antoni A and Antoni B regions. The remaining non-significant population abundances by histopathology can be found in supplementary Fig 2. **E:** Comparison of alternatively-activated TAMs versus classically-activated TAMs across both Antoni A and Antoni B regions. **F:** Relative abundance of cell populations across ROIs, annotated by case, histopathology, and disease severity. SMA: smooth muscle actin. *=p<0.05, **=p<0.01, ***=p<0.001. Statistical comparisons were made using unpaired t-tests (for normally distributed data) or Mann Whitney U tests.

PD-L1+ Schwann cells, proliferative PD-L1+ Schwann cells, vascular endothelial cells, and hypoxic perivascular neutrophils were all found to be significantly more abundant in Antoni A regions compared to Antoni B regions (Fig 1E). There was no significant difference in the remaining cell populations across the two histomorphic niches (supplementary Fig 2). Interestingly, we also observed that classically-activated TAMs were significantly more abundant in both Antoni A and B regions compared to alternatively-activated TAMs (Fig 1E). Contrasting the relative proportion of the identified cell populations across the different case ROIs within our TMA revealed substantial intratumoral heterogeneity (with differences in abundance of cell populations in different Antoni A and B ROIs within the same tumour), as well as intertumoral heterogeneity (Fig 1F), where Schwann cell and myeloid cell populations were particularly diverse across tumour regions and between cases. Given this, we addressed whether disease severity (*NF2* pathogenic variants) contributed to the observed intertumoral heterogeneity. However, we observed no significant differences in relative cell abundances in Antoni A regions between severity groups, suggesting disease severity does not correlate with intertumoral heterogeneity (supplementary Fig 3).

### The interactions of myeloid cell populations with Schwann cell populations across histomorphic niches in *NF2* SWN-related vestibular schwannoma

Given that Schwann cells and macrophages are the predominant populations in VS (Fig 1), and the apparent importance of macrophages in controlling VS growth rate^23,25^, we next investigated how Schwann cell and myeloid cell populations compartmentalize and interact within the distinct Antoni A and B niches. Representative images of canonical Schwann cell (S100B, SOX-10), myeloid cell (Iba1, HLA-DR) and stromal markers (SMA), and the spatial positioning of identified Schwann cell, myeloid, and stromal populations within Antoni A and B regions are shown in Fig 2A.

**Figure 2:**
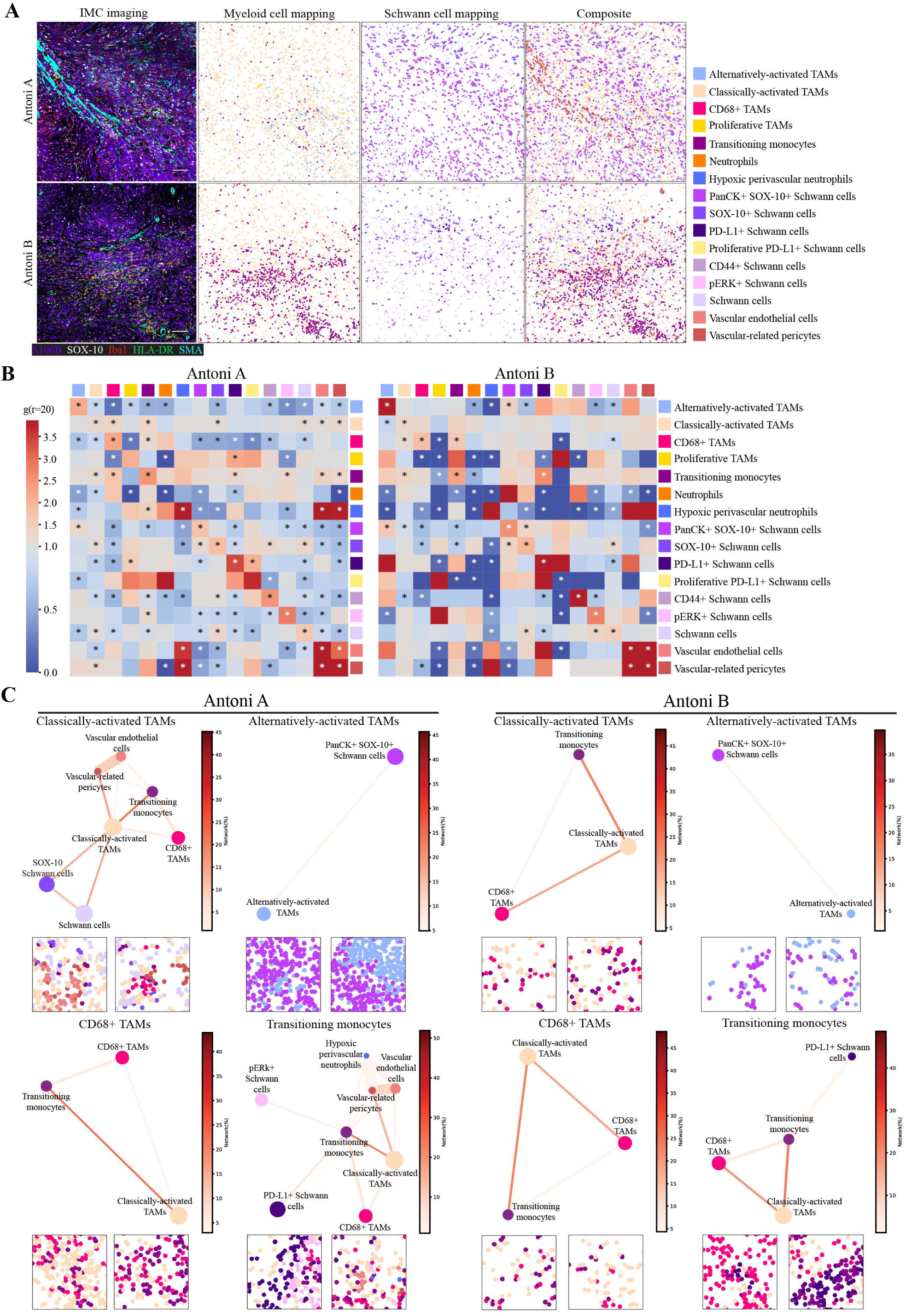
Spatial omics highlights the interactomes between Schwann cell and Myeloid cell populations in *NF2* SWN-related vestibular schwannoma. **A:** From left to right, IMC visualisation of Schwann cells and myeloid cells across Antoni A and Antoni B regions. Markers used: S100B (purple), SOX-10 (white), Iba1 (red), HLA-DR (green), and SMA (cyan). Next, single cell spatial seaborn maps of all Schwann cell and myeloid cell populations within Antoni A and B regions are shown, with a composite of the 2 major cell groups with the vascular networks. Scale bar indicates 100µm. **B:** Cross-pair correlation functions (PCF) heatmaps showing the Schwann cell and myeloid cell interaction; scale bar indicates strength of significant cell pair correlates. **C:** Spatial connectivity plots indicating the significant networks of Schwann cell and myeloid cell populations, by adjacency cell network (ACN) analysis, across Antoni A and Antoni B histomorphic niches, with representative images of each network. Node size (coloured circle) indicates mean abundance for each cell cluster across all ROIs; lines connective each node shows significant cell networks between cell types informed by CAN analysis. Line thickness associates the gr20 value to each significant cell pair, where the thicker the line, the higher the co-localisation of the cell populations. scale bar indicates strength of connectivity. **B,C:** *=p<0.05, using cross-PCF and ACN (see Methods).

To quantitatively assess the identity of and extent of homotypic (between the same cell populations) and heterotypic (between different cell populations) cell-cell interactions between myeloid cell and Schwann cell populations across the histomorphic niches, we performed cross pair correlation functions (cross-PCF) analyses^41^. In Antoni A regions (Fig 2B), myeloid and Schwann cell populations largely clustered with cells of the same type or with cells within the same sub-population group (i.e., myeloid cell populations with myeloid cells and Schwann cell populations with Schwann cells), and cells generally showed lower than expected (as if by chance) interactions with cell populations of different groups. In contrast, there were fewer statistically significant cell-cell interaction partners within Antoni B regions, with cell populations again showing preferential homotypic interactions with themselves (Fig 2B). The significant interactions between hypoxic perivascular neutrophils, neutrophils, and classically-activated TAMs was lost within the perivascular niche of Antoni B regions. Together, these data indicate there is substantial spatial variation in cell positioning and interactions between Antoni A and B regions. Whilst it appears myeloid cells and Schwann cells have a preference to interact more with themselves across the histomorphic niches, there is more cell-cell interactivity and heterogeneity between myeloid cells and Schwann cells in Antoni A regions compared to Antoni B regions. The cellular landscape and cell-cell interactions within the perivascular niche also appears to differ across regions.

To expand upon the direct cell-cell interactivity analyses, we next investigated the cellular networks of myeloid cell populations with Schwann cell populations across Antoni A and B histomorphic niches, using adjacency cell network (ACN) analysis (Fig 2C). As suggested from the cell-cell interaction analyses (Fig 2B), classically-activated TAMs and transitioning monocytes had the most heterogenous networks in Antoni A regions, with significant connectively with various Schwann cell and myeloid cell populations as well as vascular-related cells. These extensive networks were lost in Antoni B regions, where classically-activated TAMs and transitioning monocytes only connected with each other and CD68+ TAMs, with transitioning monocytes also maintaining their relation to PD-L1+ Schwann cells. Interestingly, alternatively-activated TAMs and CD68+ TAMs had the same networks in both Antoni A and B regions. Together, these analyses indicate distinct cellular networks that are either heterogenous (as with classically activated TAMs and transitioning monocytes) or homogenous (as with alternatively-activated TAMs and CD68+ TAMs) between myeloid cells in Antoni A regions, which are lost (heterogeneous) or maintained (homogenous) in Antoni B regions.

### T-cell populations exist within distinct environments associated with infiltration and suppression across histomorphic niches in *NF2* SWN-related vestibular schwannoma

T-cells play a fundamental role in anti-tumoral immunity and tumour cell elimination^43^, yet their role in VS is unclear. We identified different populations of CD8+ and CD4+ T-cells within Antoni A and B regions (Fig 1D), but what these populations interact with and if they occupy the same tumour subniches is unknown. As such, we investigated the spatial interactions of T-cells across the different histomorphic Antoni A and B niches in *NF2* SWN VS. Representative images of canonical T-cell (CD8, CD4), Schwann cell (S100B), macrophage (Iba1) and stromal (SMA) marker staining, and the spatial positioning of identified T-cell, Schwann cell, myeloid, and stromal populations within Antoni A and B regions are shown in Figure 3A.

**Figure 3:**
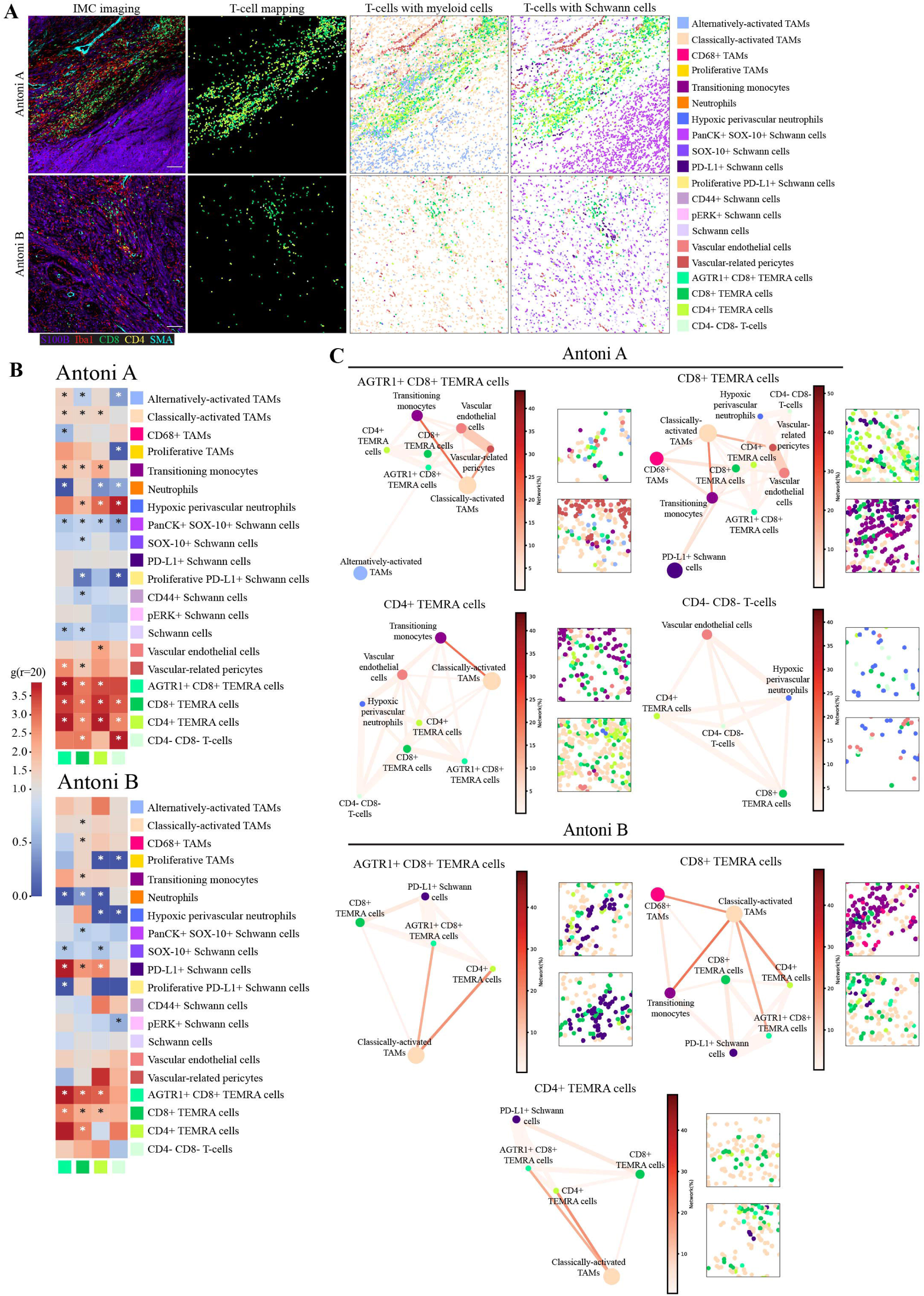
Spatial omics demonstrates the T-cell interactome across different histomorphic niches in *NF2* SWN-related vestibular schwannoma. **A:** From left to right, IMC visualisation of T-cell across Antoni A and Antoni B regions. Markers used: S100B (purple), Iba1 (red), CD8α (green), CD4 (white) and SMA (cyan). Next, single cell spatial seaborn maps of all T-cell populations, and co-localisation to Schwann cell or TAM populations with the vascular networks across Antoni A and B regions. Scale bar indicates 10. **B:** Cross-pair correlation functions (PCF) heatmaps showing T-cell interaction partners with all other cell populations; scale bar indicates strength of significant cell pair correlates. **C:** Spatial connectivity plots indicating the significant networks of T-cell populations, by adjacency cell network (ACN) analysis, across Antoni A and Antoni B histomorphic niches, with representative images of each network. Node size (coloured circle) indicates mean abundance for each cell cluster across all ROIs; lines connective each node shows significant cell networks between cell types informed by CAN analysis. Line thickness associates the gr20 value to each significant cell pair, where the thicker the line, the higher the co-localisation of the cell populations. Scale bar indicates strength of connectivity. **B,C:** *=p<0.05, using cross-PCF and ACN (see Methods).

To statistically evaluate the direct cell-cell interactome of the four identified T-cell populations across histomorphic niches in VS tumours, we again employed cross-PCF analysis^41^. In Antoni A regions (Fig 3B), the strongest interactions were seen between T-cell subtypes, with AGTR1+ CD8+ T effector memory re-expressing CD45RA (TEMRA) cells, CD8+ TEMRA cells, CD4+ TEMRA cells, and CD4-CD8-T-cells all appearing to interact with themselves and each other. This indicates, as shown in Figure 3A, that the different T-cell populations frequently intermix in Antoni A regions, rather than segregate within distinct areas. Interestingly, both CD8+ TEMRA populations interacted with vascular-related pericytes, whilst CD4+ TEMRA cells clustered with vascular endothelial cells. Both CD8+ TEMRA populations and CD4+ TERMA cells had significant interactions with transitioning monocytes and classically-activated TAMs, but only AGTR1+ CD8+ TEMRA cells significantly engaged with alternatively-activated TAMs. Conversely, all T-cell populations except for AGTR1+ TEMRA cells significantly interacted with hypoxic perivascular neutrophils. As observed in Figure 3A, the T-cell populations showed lower than expected interactions with all the different Schwann cell populations, suggesting T-cells are predominantly sequestered within TAM-rich areas in Antoni A regions of the tumour. Interestingly, there were, in general, fewer clustering interactions across T-cell populations in Antoni B regions (Fig 3B). AGTR1+ TEMRA cells and CD8+ TEMRA cells still exhibited significant homotypic and heterotypic interactions with themselves and each other, as well as with CD4+ TEMRA cells. On the other hand, CD4-CD8-T-cells and CD4+ TEMRA cells exhibited lower statistical cell-cell interactions with other T-cell populations within Antoni B regions than observed in Antoni A regions. As opposed to the results in Antoni A regions, neither CD8+ TEMRA cells or CD4+ TEMRA cells interacted significantly with vascular-related pericytes or vascular endothelial cells in Antoni B regions, and only CD8+ TEMRA cells showed interactions with classically-activated TAMs or transitioning monocytes. Instead, the CD8+ TEMRA cells and CD4+ TEMRA cells showed statistically enriched interactions with PD-L1+ Schwann cells within Antoni B regions. Taken together, these results highlight the heterogeneity of T-cell interactions across different histomorphic niches in VS, particularly when addressing T-cell co-localisation with TAM populations and immunoregulatory PD-L1+ Schwann cells in Antoni A and B regions, respectively.

Given the apparent differences in cell-cell interactions and positioning of T-cell populations in Antoni A and B regions, we next investigated the cellular networks of T-cell populations across these histomorphic niches by ACN analysis (Fig 3C). Across both histomorphic niches, T-cells commonly had a strong network with each other, particularly in Antoni A regions. AGTR1+ CD8+ TEMRA cells had connectivity with both classically-activated and alternatively-activated TAMs, transitioning monocytes and vascular-related cells in Antoni A regions, whereas in Antoni B regions, these T-cells lost their network with vasculature and alternatively-activated TAMs, but gained significant connectivity to PD-L1+ Schwann cells. CD8+ TEMRA cells had the most heterogeneous networks across both histomorphic niches, particularly in Antoni A regions, with connections to vascular-related cells, classically-activated TAMs, transitioning monocytes, CD68+ TAMs and PD-L1+ Schwann cells. These networks were altered in Antoni B regions, where CD8+ TEMRA cells lost connectivity to the vascular network, most myeloid cells, and showed high connectivity with PD-L1+ Schwann cells. CD4+ TEMRA cells also lost their connectivity to vasculature (and transitioning monocytes) from Antoni A to Antoni B regions, and gained significant connections to PD-L1+ Schwann cells, whilst maintaining their connection to classically-activated TAMs. Interestingly, the network between double negative CD4-CD8-T-cells and other T-cell populations and vasculature in Antoni A regions was completely lost in Antoni B regions, with no significant network connections present. Together, these analyses suggest T-cells in Antoni A regions exist within networks associated with infiltration and localisation with myeloid cells, whilst in Antoni B regions, these networks completely alter and T-cells become significantly more associated with immunoregulatory PD-L1+ Schwann cells, highlighting distinct modes of T-cell regulation that are niche-dependent.

### Derivation of cellular neighborhoods reveals distinctive spatial organisation and immune environments across histomorphic niches in *NF2* SWN-related vestibular schwannoma

To further interrogate the topography of the Antoni A and B TME, we deconstructed the tumour ROIs into cellular neighbourhoods (CNs). CNs are collectively defined by both the identity and proportionality of clustering cells and tumour stroma within a tissue space, and the derivation of CNs allows for visualization of distinct sub-niches within heterogeneous tissues, such as tumours. To this end, we quantified CNs using CellCharter^42^. We initially computed CNs across all ROIs, assimilating both Antoni A and B regions. Whilst this identified a statistically optimal 14 distinct CNs (supplementary Fig 4A and B), due to the increased number of Antoni A compared to Antoni B ROIs within our dataset, these CNs were dominantly distributed within Antoni A regions; the discrete cellular networks identified in Antoni B ROIs (Figures 2 and 3) were lost within this combined CN analysis (supplementary Fig 4C). Hence, we generated unique CN profiles for both Antoni A and B regions independent of one another. In this instance, the optimal stability was achieved at *n*=10 clusters for both Antoni A and B regions (supplementary Fig 4D and E).

In Antoni A regions (Fig 4A), we identified CNs relating to inflammatory schwannoma niches characterised by pERK and PD-L1 expression (CNs A1, A5, and A7), injury response-like schwannoma niches with co-localisation of alternatively-activated TAMs and PanCK+SOX-10+ Schwann cells (CNs A0 and A2), myeloid-rich niches (CNs A4), and T-cell rich niches illustrative of perivascular infiltration and co-localisation with TAMs (CNs A3, A6 and A8). CN A9 was primarily comprised of ‘Other’ cell populations, and some Schwann cells. Interestingly, some of these Antoni A CNs were mirrored in the independently-derived Antoni B CNs, including inflammatory schwannoma niches (CN B8), injury response-like schwannoma niches (CNs B4 and B7), myeloid-rich niches (CN B6), and immune-rich perivascular niches (CN B2 and B9). We also identified niches unique to Antoni B regions, namely demonstrating T-cell regulation through PD-L1 and alternatively-activated TAMs, as well as TAM proliferation (CNs B0 and B3). CNs B1 and B5 were largely comprised of the ‘Other’ cell populations. We also quantified the abundance of CNs across cases (Fig 4B), which reflected the single-cell composition and heterogeneity seen in Fig 1G.

**Figure 4:**
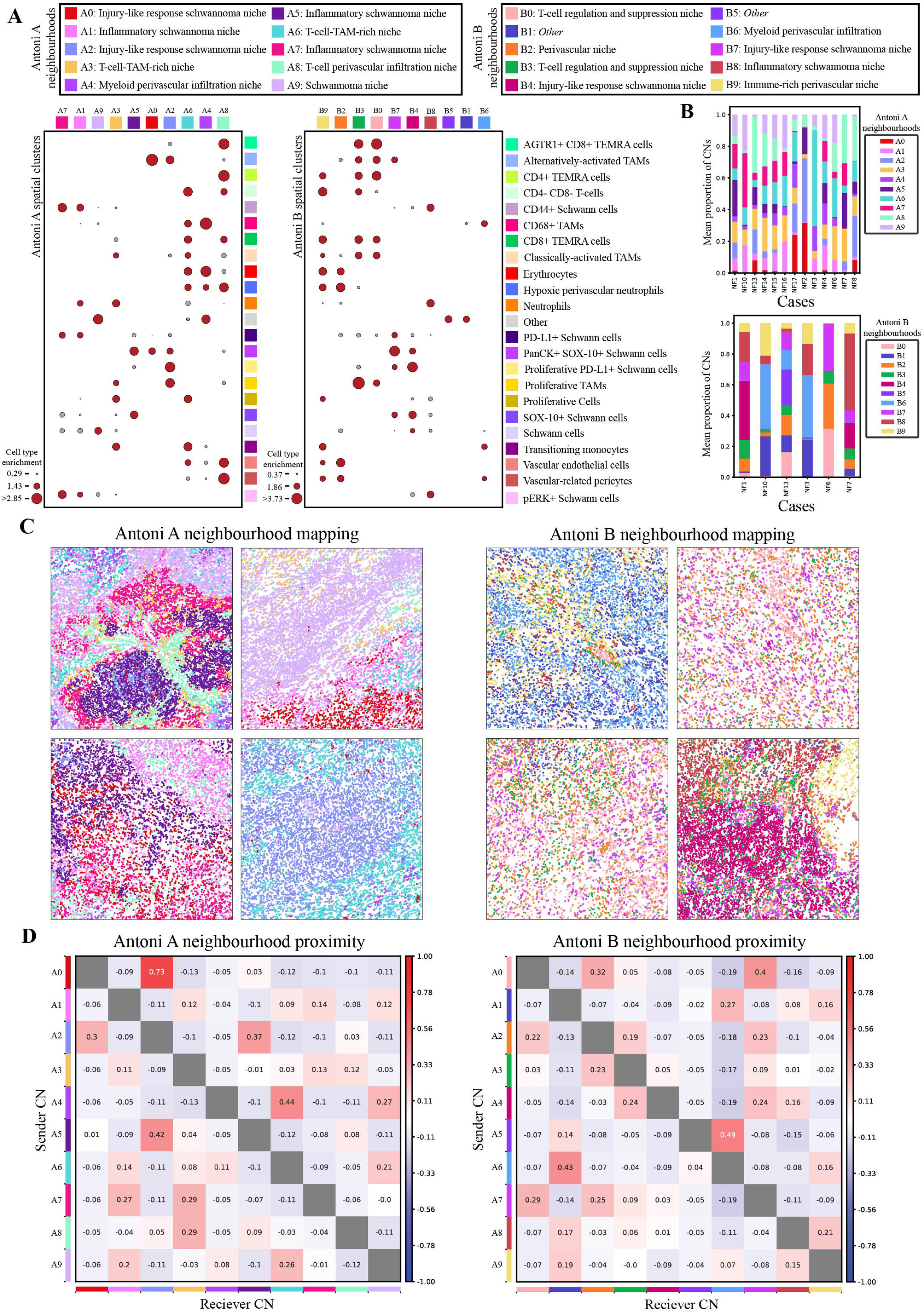
Cellular neighbourhood analysis identifies discrete microenvironmental niches across histomorphic niches in *NF2* SWN-related vestibular schwannoma. **A:** Dot plot of the individually derived spatial clusters (referred to as cellular neighbourhoods, CN) for Antoni A regions and Antoni B regions. Optimal cluster stability for both groups was 10. Cells were clustered using Gaussian mixture modelling. Node size indicates relative cell type enrichment of each cell across generated CNs (red = high enrichment, grey = low enrichment). **B:** Relative abundance of both Antoni A and Antoni B CNs across cases. **C:** Representative spatial mapping of CNs across Antoni A and Antoni B regions. Each image is representative on an independent case to highlight intertumoral heterogeneity. **D:** CN proximity heatmaps depicting the spatial proximity of each CN to every other CN. Y-axis indicates sender communication; x-axis indicates receiver communication. Diverging scale bar denotes proximity enrichment, with red specifying high proximity, and blue specifying low proximity.

Next, we investigated the spatial layout of the different CNs in Antoni A and B regions. For the most part, Antoni A regions were comprised of similar CN densities relating to the different biological processes outlined above and appeared to occupy distinct tissue regions as shown in the representative images in Fig 4C. This would suggest that Antoni A regions had a high level of organisation. In contrast, Antoni B regions had more varied CN profiles, where some CNs (particularly CNs B4, B6 and B8) dominated the cellular landscape of these regions. Spatially, Antoni B CNs were more inter-dispersed across the tissue regions (Fig 4C) compared to Antoni A CNs, indicating there is less organisation of CNs in Antoni B regions compared to Antoni A regions. This is particularly true for CNs B0 and B3, suggesting that T-cell regulation in these regions occurs through discrete interactions throughout the tissue as opposed to specific focal aggregates.

Finally, we assessed the neighbourhood proximity between CNs (Fig 4D) within the two histomorphic niches, as communication between neighbourhoods likely influences biological functions within the TME. In both Antoni A and B regions, it appeared that CNs of similar compositon were spatially close with each other. CN A0 had the strongest proximity to CN A2 in Antoni A regions, both of which exhibited injury response-like processes. The inflammatory tumour niches (CNs A1, A5 and A7) were closely associated with immune-rich niches (namely CNs A3, A6 and A8), as well as other inflammatory tumour niches, suggesting an inflammation-driven recruitment of immune cells within these regions. In Antoni B regions, we saw similar preferential proximity between the injury response-like niches (CNs B4 and B7), with these niches also proximally associating with niches involving T-cell suppression and TAM proliferation (CN B0 and B3). The latter CNs were also enriched for alternatively-activated TAMs, whereby these TAMs likely support responses involving wound repair. T-cells in Antoni B regions appeared to cluster within these suppressive niches, however, some resided within CN B9 and proximally co-localised with CN B6 perivascular niches. Collectively, our analyses suggest that Antoni A and B regions demonstrate a degree of spatial organisation with CNs associated with inflammation and injury-like responses conserved across both regions, however, Antoni B regions appear less organised than Antoni A regions, which may drive formation of T-cell suppressive niches across these tissue regions.

### Bevacizumab treatment significantly increases the expression of CD44 on Schwann cells, and reduces AGTR1 expression on T-cells, in *NF2* SWN-related vestibular schwannoma

VS tumours in individuals with *NF2* SWN are heterogeneous both clinically and therapeutically, and as we have demonstrated this heterogeneity extends to the histopathological, cellular and spatial levels, highlighting potential for improved patient stratification and better management of the disease. To this end, we investigated the effects of Bevacizumab treatment on the TME of VS from 3 patients where bevacizumab failed to control tumour growth, to enable identification of potential mechanisms leading to bevacizumab failure, and alternative modes of therapeutic intervention. These 3 cases were from *NF2* SWN patients where bevacizumab treatment was stopped prior to resection (outlined in Table 1). For this analysis, we focused only on Antoni A ROIs, as Antoni B ROIs could not be represented in the bevacizumab-treated group.

Representative images of canonical Schwann cell, macrophage, T-cell, and stromal cell markers and the spatial mapping of identified cell populations between treatment naïve and bevacizumab-treated cases are shown in Fig 5A. Interestingly, CD44+ Schwann cells were significantly more abundant, but AGTR1+ CD8+ TEMRA cells were significantly less abundant, in bevacizumab-treated cases compared to treatment-naïve cases (Fig 5B). No significant differences were seen between the other cells types between treatment-naïve and bevacizumab-treated groups (supplementary Fig 5A). We also compared the ratio of classically-activated TAMs to alternatively-activated TAMs between the two groups, and found that alternatively-activated TAMs were significantly less abundant in both treatment-naïve and bevacizumab-treated cases (Fig 5C). We next compared the CN composition between treatment naïve and bevacizumab-treated cases, mapped in Fig 5D; we did this by re-generating the Antoni A CNs with the additional bevacizumab cases included. The CNs from the combined naïve and bevacizumab analysis were very similar to the naïve-only CNs (supplementary Fig 6B), characterised by microenvironmental processes relating to inflammation, injury-like responses, and immune enrichment and infiltration (supplementary Fig 5B). When assessing the proportionality of CNs between naïve and bevacizumab cases (Fig 5E), CNs Bev-0, 1 and 9 were noticeable different, with CN Bev-9 being significantly more abundant in bevacizumab than naïve cases (Fig 5E). This CN is enriched for CD44+ Schwann cells, which parallels the significant increase in CD44+ Schwann cells in bevacizumab-treated cases (Fig 5B). Taken together, these analyses suggest that bevacizumab-treated tumours from patients who experienced bevacizumab failure are enriched for Schwann cell populations that may indicate ECM-related remodeling processes, as well as potential changes to the immune landscape of VS.

**Figure 5:**
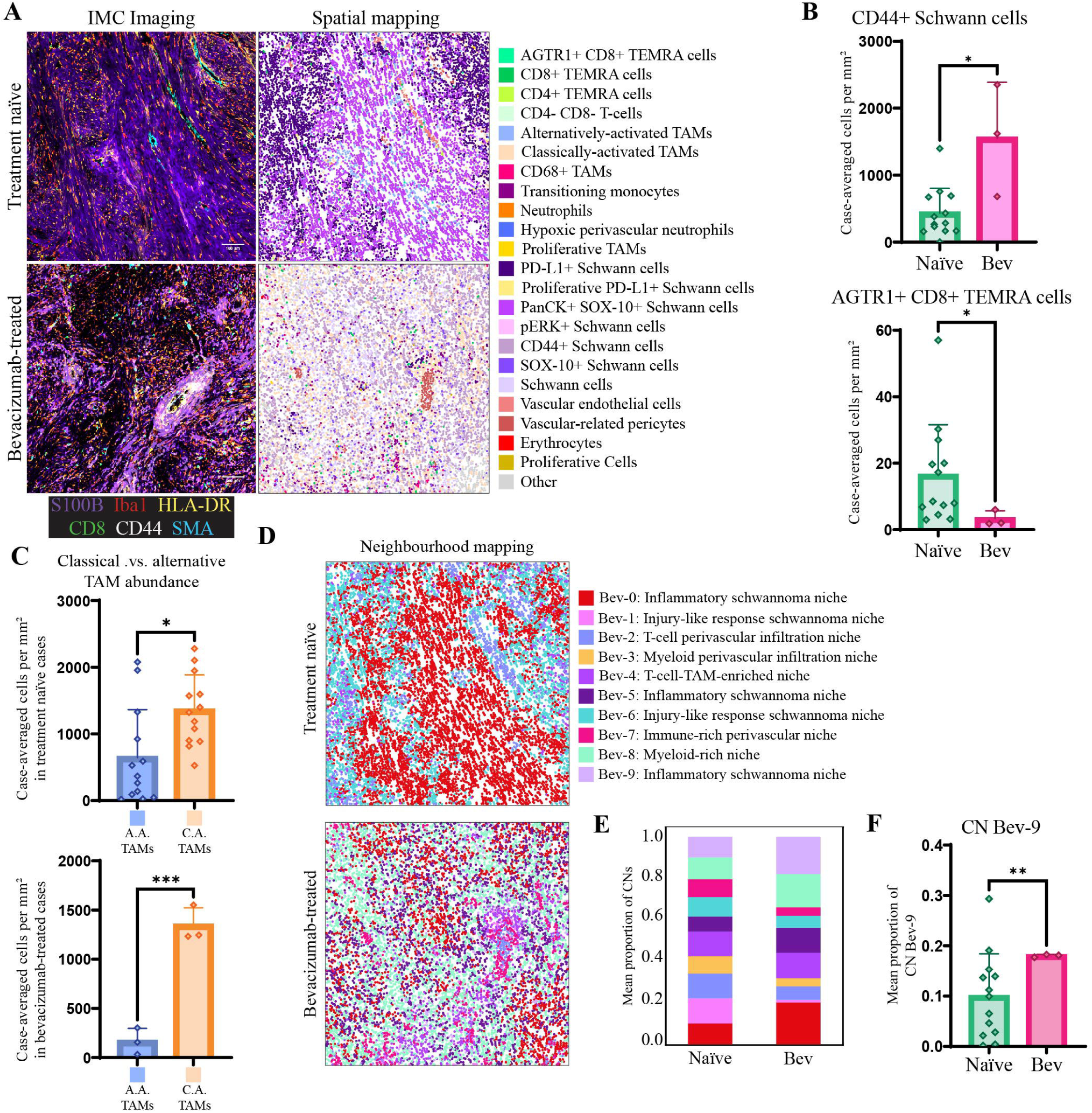
IMC analyses highlight the influence of bevacizumab treatment on the microenvironmental in *NF2* SWN-related vestibular schwannoma. **A:** From left to right, IMC visualisation of core tumour and immune-related markers across treatment naïve and bevacizumab-treated groups. Markers used: S100B (purple), Iba1 (red), CD8α (green), CD44 (white), HLA-DR (yellow) and SMA (cyan). Spatial seaborn maps of all populations across treatment naïve and bevacizumab-treated groups. Scale bar indicates 100µm**. B:** Abundance of significantly different cell populations across treatment naïve and bevacizumab-treated groups, tested by Mann Whitney U test. The remaining non-significant population abundances by treatment status can be found in supplementary Fig 6. **C:** Comparison of alternatively-activated TAMs versus classically-activated TAMs across treatment naïve and bevacizumab-treated groups. Treatment-naïve tested by Mann Whiteney U test, and bevacizumab-treated tested by unpaired t-test. **D:** Visualisation of bevacizumab-inclusive CNs across treatment naïve and bevacizumab-treated groups. Dot plot of spatial clusters can be found in supplementary Fig 6B. E: Mean proportion of CNs across treatment-naïve and bevacizumab treated groups. **F:** Comparison of CN Bev-9 between treatment-naïve and bevacizumab treated groups, tested by Welch’s t-test. *=p<0.05, **=p<0.01, ***=p<0.001. Bev = bevacizumab.

## Discussion

In this study we have employed high dimensional imaging to analyse in detail the spatial architecture of *NF2* SWN-related VS, highlighting that these tumours exhibit a complex topology with tumour-rich and immune cell-rich zones across histomorphic niche Antoni A and B regions.

By IMC analyses, we identified 23 distinct cell populations relating to different phenotypes and activation states of Schwann cells, myeloid cells, and lymphocytes, as well as tumour stroma, across the histopathological landscape of VS. This further reinforces the complexity and cellular heterogeneity of these tumours. It is well established that TAMs, along with Schwann cells, are the most abundant cell type within both sporadic and *NF2* SWN-related VS^27,44^. Our study indicates that classically-activated pro-inflammatory TAMs make up a greater proportion of the myeloid compartment than alternatively-activated TAMs in *NF2* SWN-related VS, when calculating the relative ratio of abundance of the two myeloid sub-populations across both Antoni A and B regions. The inflammatory mediators secreted from the TME likely drives recruitment of monocytes into the tumour and support the TAM-rich identity of VS. Indeed, previous immune profiling studies in VS have shown that schwannoma cells and TAMs secrete inflammatory factors, such as interleukin (IL)-2, IL-6 and CCL20, and that increased levels of circulating monocytic chemokines are observed in plasma samples from growing VS cases^45,46^. Supporting this, we have shown that classically-activated TAMs and transitioning monocytes have extensive interaction networks in Antoni A regions, namely with other myeloid cells, vasculature and some Schwann cell populations. Furthermore, our neighbourhood analyses demonstrated the existence of prominent perivascular networks that co-localise with inflammatory tumour regions. As such, our data suggests that there is active recruitment of monocytes through the perivascular niche in VS, which transition into classically-activated TAMs and promote tumour inflammation in Antoni A regions. This poses the question as to whether VS growth is driven by inflammation or clonal expansion of neoplastic Schwann cells, or potentially both?

The direct cell interactions between classically-activated TAMs and transitioning monocytes with vascular-related cells appeared lost in Antoni B regions within our analyses. The apparent disruption to direct cell-cell interactivity within the perivascular niche of Antoni B regions may be a consequence of degenerative or pathological changes to tumour vasculature, such as hyalinization^47^. Antoni B regions have previously been proposed to result from degeneration of Antoni A regions, given their pathological similarities to Wallerian retrograde degeneration seen in traumatic nerve injury^48^. Together with our data, this suggests that the incidence of these distinct histomorphic niches may related to spatial changes in inflammatory responses, and potentially ECM deposition, within the TME of VS.

The concept that Schwannomas arise as a consequence of injurious stimuli to peripheral nerves is evidenced by recent single-cell transcriptomic and epigenetic studies where nerve injury-like states were discovered, and characterised by myeloid cell infiltration^49,50^. Our analyses further support this finding, whereby we illustrate that alternatively-activated TAMs significantly interact with PanCK+ SOX-10+ Schwann cells in both Antoni A and B regions; cell populations expressing both PanCK and vimentin (which is present on all Schwann cell populations we identified) are associated with wound healing responses^51^, which mirror the function of alternatively-activated TAMs. Our neighbourhood analyses also highlight these injury-like niches along with PD-L1+ Schwann cells across both Antoni A and B regions. Taken together, this suggests that injury-like niches in VS are conserved across histomorphic regions, and likely support the polarisation and maintenance of alternatively-activated TAMs in a classically-activated TAM-dominated landscape.

Given the importance of T-cells in other tumour types as key immunotherapeutic targets^52,53^, we also interrogated the positioning and proximity of both CD4+ and CD8+ T-cells within *NF2* SWN-related VS. One subset of TEMRA cells expressed angiotensin II type 1 receptor (AGTR1), which has been implicated in the regulation of T-cell expansion, differentiation and function during *Plasmodium* infection^54^; the role of AGTR1 on CD8+ T-cells in VS remains to be tested. The identified T-cells exhibited unique spatial interactions and networks between Antoni A and B regions in VS. T-cells within Antoni A regions strongly interacted with other T-cells and myeloid cells (in particular classically-activated TAMs and transitioning monocytes), as well as vasculature, but did not appear to interact with Schwann cells. This indicates that T-cells may be sequestered in the TAM-enriched areas in Antoni A regions and are unable to directly target tumour cells. It has been noted that TAMs can form long-lasting interactions with T-cells and impede their ability to interact with tumour cells, which directly limits the efficacy of anti-PD-1 therapies^55^. It is likely that secretion of chemokines by classically-activated TAMs is drives the significant co-localisation between T-cells and TAMs in VS, establishing a T-cell exclusionary niche in Antoni A regions^56^. Interestingly, we observed that T-cell positioning was altered within Antoni B regions, where some T-cells predominantly interacted with PD-L1+ Schwann cells. Furthermore, our neighbourhood analyses illustrate that CD8+ and CD4+ T-cells exist within niches with PD-L1+ Schwann cells and alternatively-activated TAMs in Antoni B regions, both of which are detrimental to T-cell functionality. A better understanding of how these spatially-dependent responses are orchestrated and maintained in VS could be leveraged to enable maximal T-cell-mediated tumour killing, through the potential disruption of TAM sequestration in Antoni A regions, and the blockade of inhibitory signaling provided by PD-1-PD-L1 interaction in Antoni B regions. As such, our spatial analyses highlight a clinical rationale for selected combinatorial treatment for VS tumours in *NF2* SWN.

Finally, we investigated the effects of Bevacizumab treatment on the TME of *NF2* SWN-related VS. Whilst Bevacizumab is not approved for *NF2* SWN by drug regulation authorities such as the FDA or MHRA, it is utilised off-label and has shown success in some *NF2* SWN patients^16,17^, and is approved for treatment by NHS England in the highly specialized commissioned NF2 service. Hence, it is important to understand why Bevacizumab treatment might fail, and if alternative pathways are upregulated in Bevacizumab-treated tumours that could be targeted therapeutically. Our results suggest that Bevacizumab treatment may selectively change the ratio of classically-activated TAMs to alternatively-activated TAMs, preferentially causing loss of alternatively-activated TAMs. Additionally, we found that Bevacizumab-treated cases had significantly more CD44+ Schwann cells, yet significantly less AGTR1+ CD8+ TEMRA cells, than treatment-naïve cases. CD44, also known as the hyaluronan receptor, has various implications for cell survival, proliferation, mobility, and is often dysregulated in cancer, leading to metastasis, fibrosis and therapy resistance^57,58^. The upregulation of CD44 on Schwann cells in Bevacizumab-treated VS cases may suggest the development of fibrotic responses, which may subsequently concur with Bevacizumab-resistance and failure. Interestingly, the reduction of AGTR1 expression on CD8+ TEMRA cells suggests that Bevacizumab may indirectly inhibit angiotensin signaling, which is a pathway targeted by the drug Losartan, which is being employed in clinical trials for VS^59,60^. Together with previous data within preclinical models of *NF2* SWN VS, may indicate Losartan as an alternative for *NF2* SWN patients experiencing Bevacizumab failure, which may be accompanied by increased matrix remodeling. However, a greater understanding of how ECM remodelling occurs in VS, and how Bevacizumab treatment changes that TME in responsive tumours, is required to prove this.

In conclusion, we present the first high dimensional deconstruction of *NF2* SWN-related VS, where we illustrate the immune landscape is associated with clear regional differences in T-cell engagement with TAMs and Schwann cells, as well as evidence of diverse pro-inflammatory and anti-inflammatory immune and Schwann cell networks and neighbourhoods. Furthermore, we suggest potential changes in the VS TME that occur following Bevacizumab treatment, and the potential link to Bevacizumab failure. Although not studied here, a key consideration will be how these vary in individuals with different symptoms, how they evolve in primary versus recurrent tumours, and how they differ in tumours with different pre-surgical growth rates. Nevertheless, our results give new insights into the tumour features and cellular pathways that may be amenable for targeting to improve treatment of *NF2* SWN-related VS.

## Acknowledgements

We thank Dr Gareth Howell and Dr Jennifer Baron within the Flow Cytometry Core Facility at the University of Manchester for helping with the imaging mass cytometry work performed in this study. We thank Dr Peter March in the Bioimaging Core Facility at the University of Manchester for helping with the microscopy work. We thank Professor Federico Roncaroli within the Department of Cellular Pathology at the Salford Royal Hospital for annotating the H &E-stained slides to identify Antoni A and B regions for incorporation within the Tissue Microarrays, which were generated by Dr Garry Ashton and team at the The Christie Hospital (CRUK), Manchester.

## Funding

The work was funded by *NF2* BioSolutions (PhD studentships supporting A.P.J and G.E.G), the Medical Research Council (MR/T016515/1 to D.B and K.N.C), and the NIHR Manchester Biomedical Research Centre (NIHR203308 to D.B and O.N.P; IS-BRC-1215-20007 to D.G.E, D.B. and O.N.P). The imaging mass cytometer used within the study was purchased through a BBSRC Alert18 award (BB/S019324/1 to K.N.C.).

## Author contributions

Conceptualization: APJ, KC, GEG

Methodology: MJH

Investigation: APJ, GEG, AKS

Resources: CJH, DGL, PO, LDB

Software: MJH, APJ

Visualization: APJ

Supervision: KC, ONP, DB, PP

Writing - original draft: APJ, KC

Writing - review & editing: APJ, KC, ONP, DB, PP, DGE, ATK, MJS, PO, DGL, CJH, MJH, GEG

Funding acquisition: KC, ONP, DB, DGE

## Competing Interests

The authors report no competing interests

## Notes

### Competing Interest Statement

The authors have declared no competing interest.

