## Supplementary figures and images for "High-Dimensional Imaging of Vestibular Schwannoma Reveals Distinctive Immunological Networks Across Histomorphic Niches in *NF2*-related Schwannomatosis"

### Supplementary Figure 1

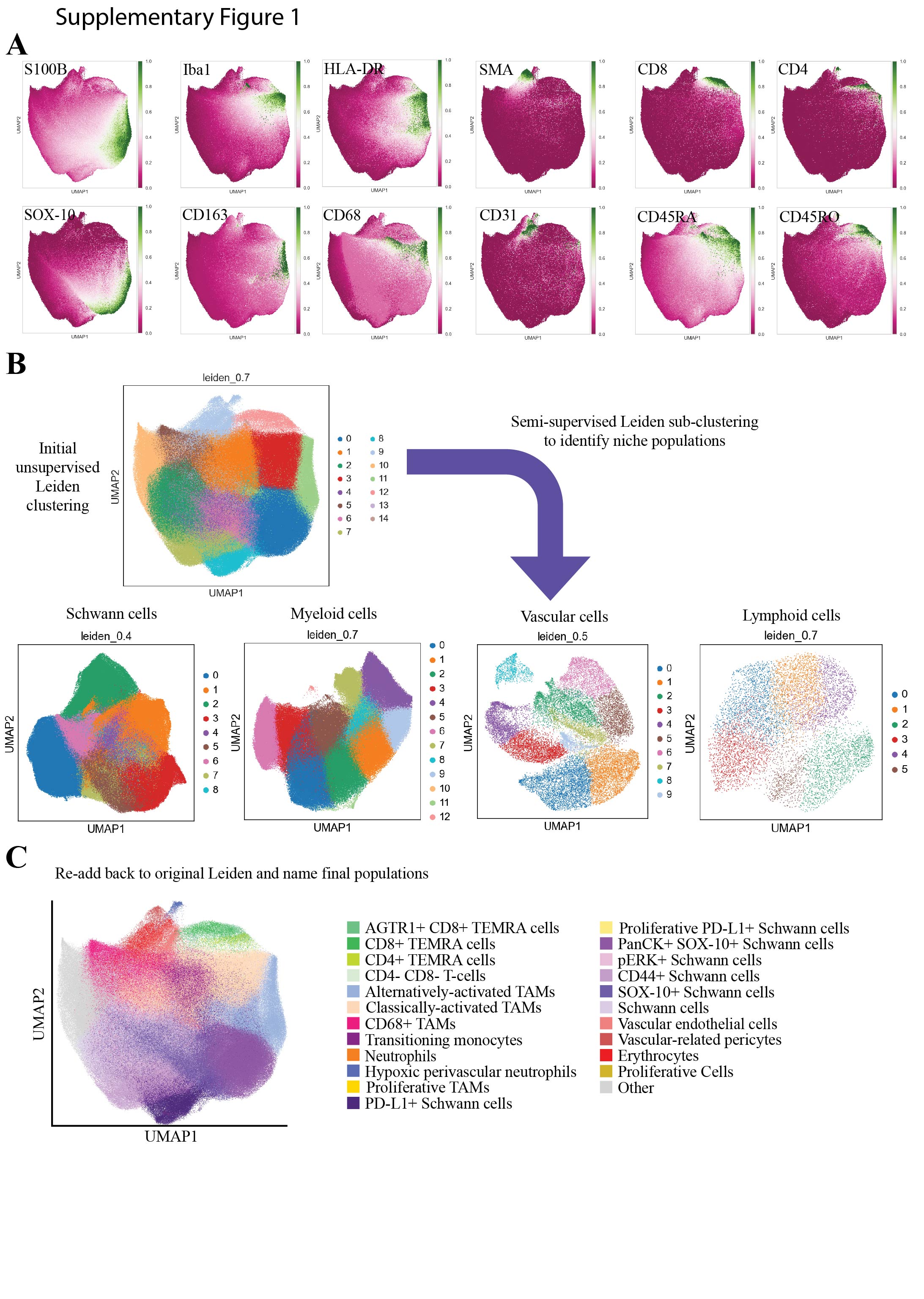

### Supplementary Figure 2

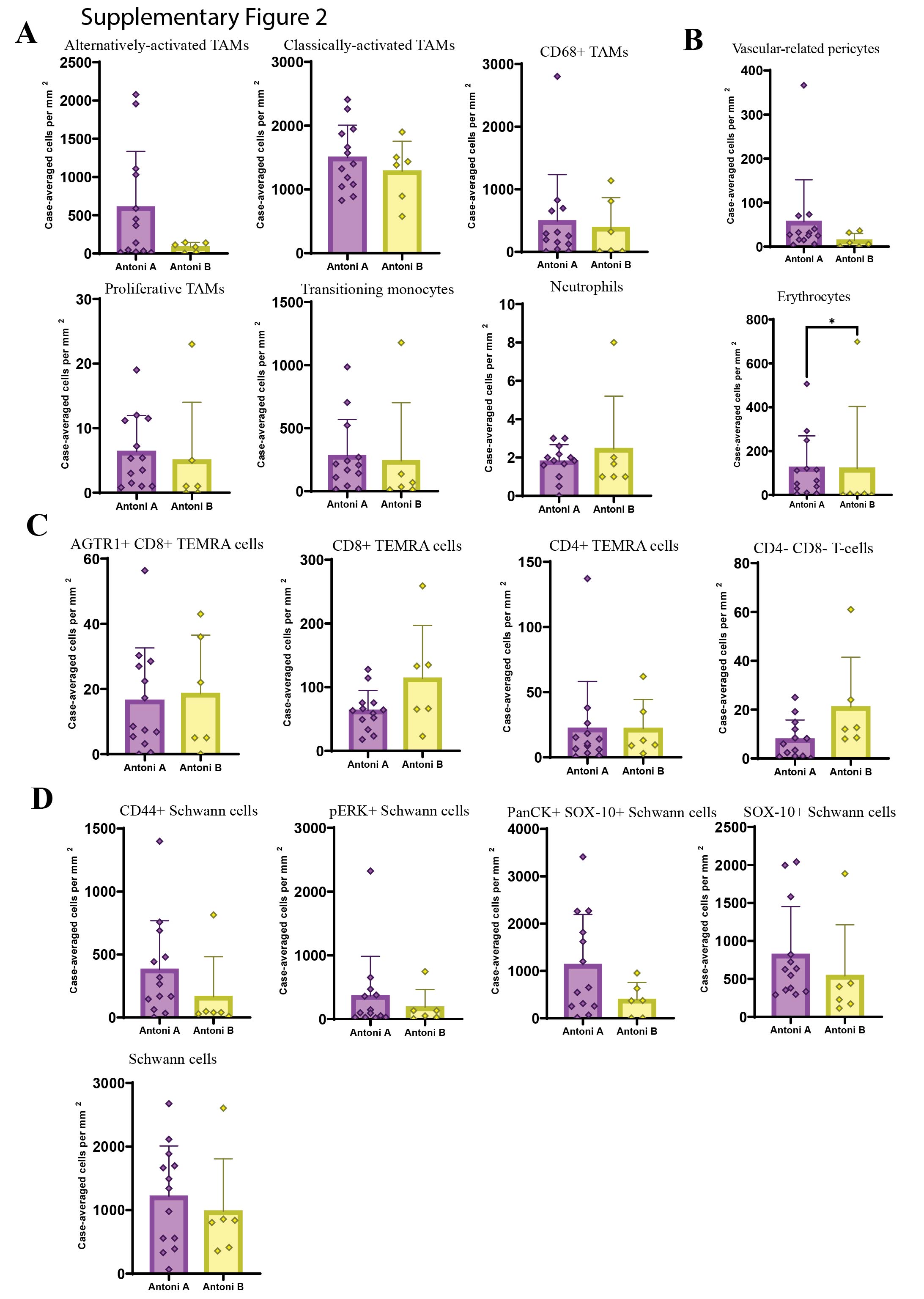

### Supplementary Figure 3

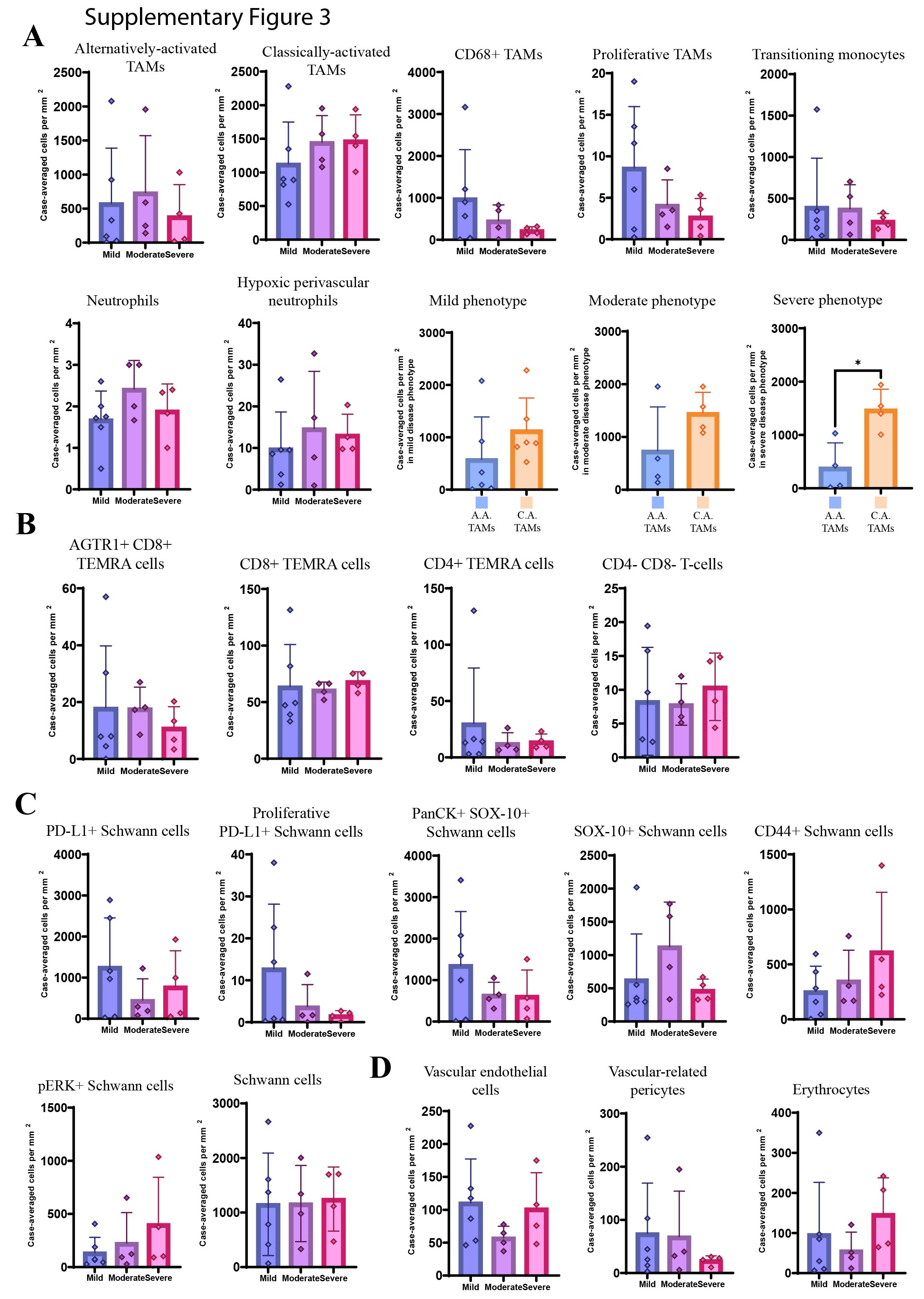

### Supplementary Figure 4

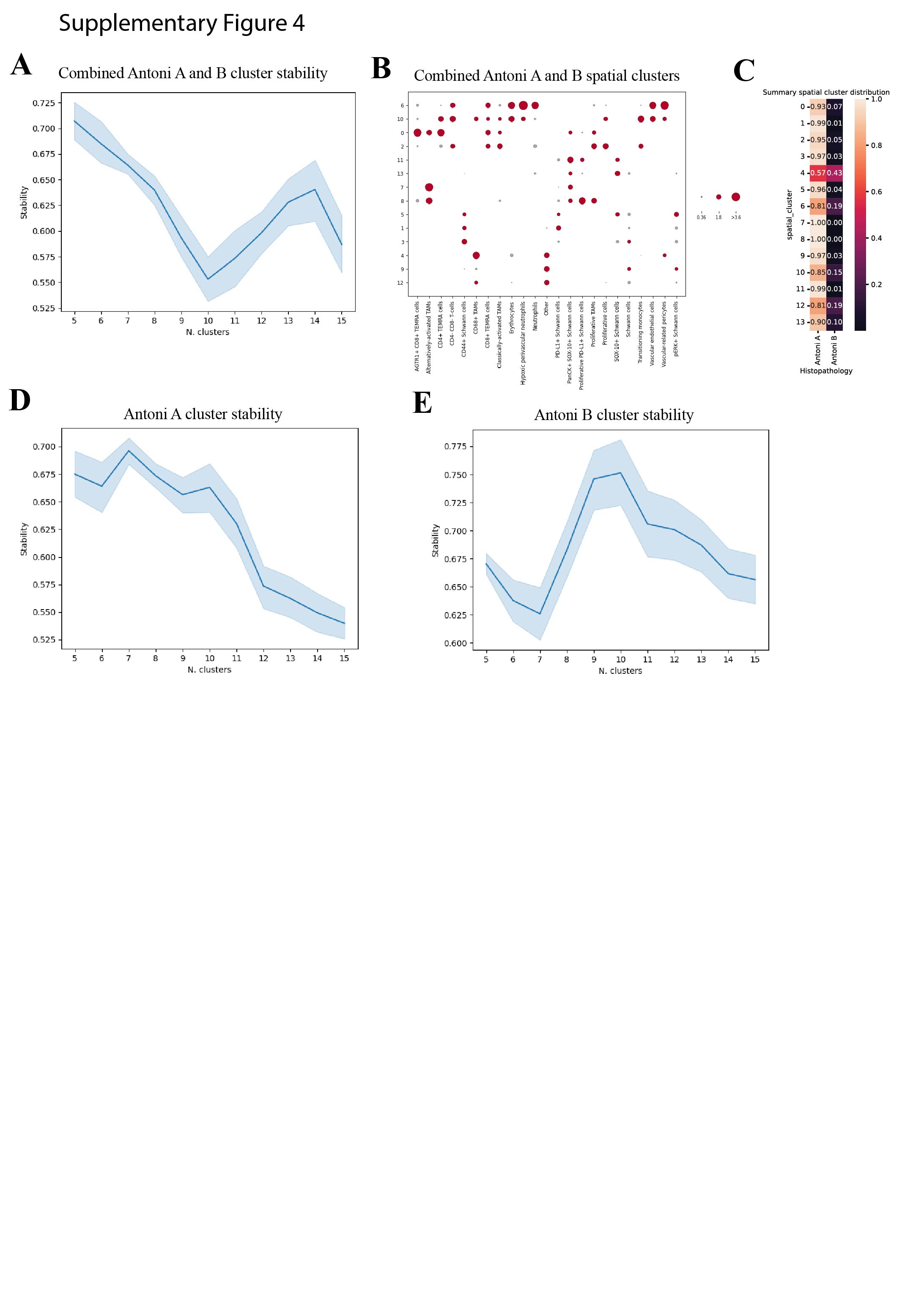

### Supplementary Figure 5

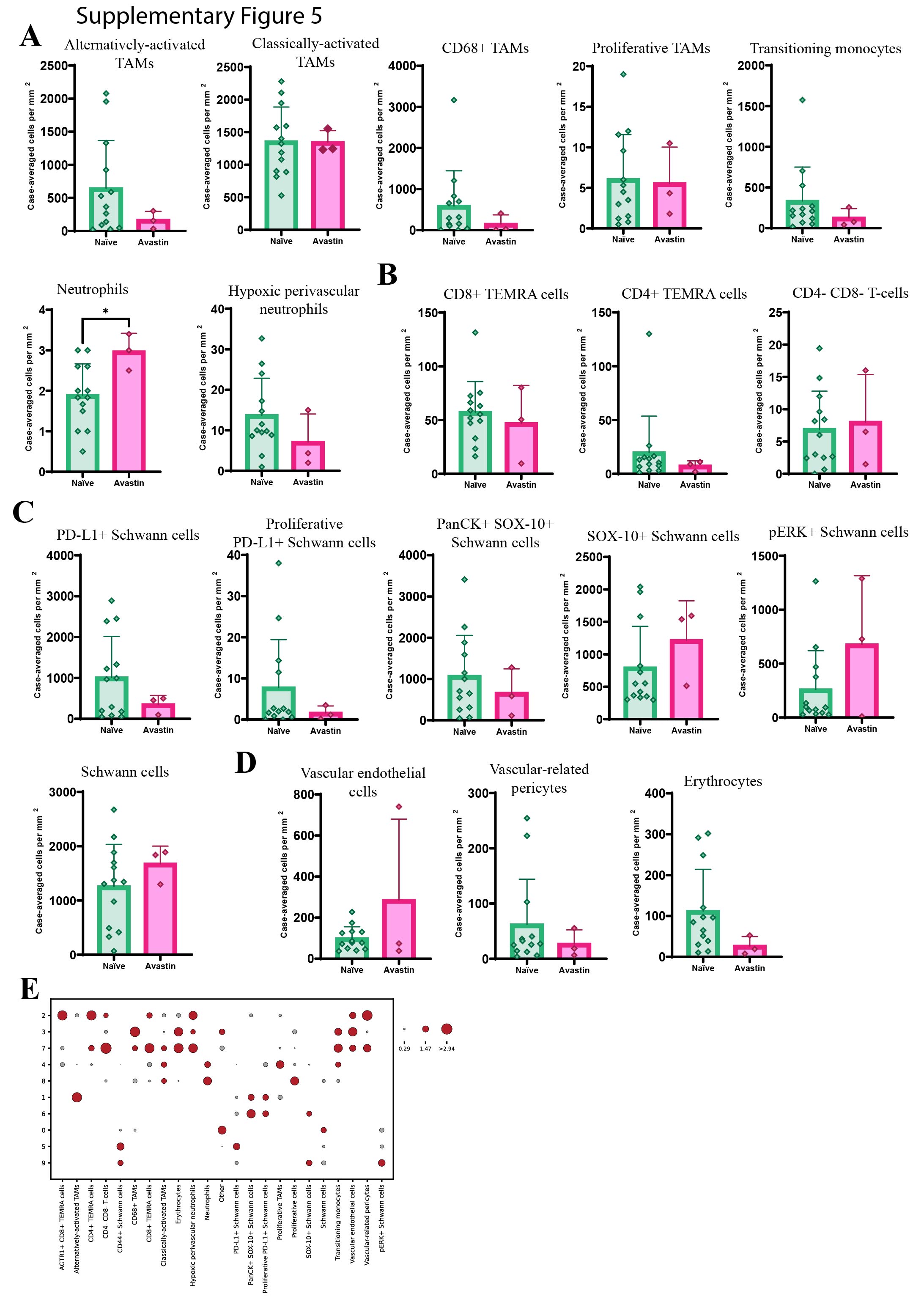
