## Supplementary Figure Legend 1 for "High-Dimensional Imaging of Vestibular Schwannoma Reveals Distinctive Immunological Networks Across Histomorphic Niches in *NF2*-related Schwannomatosis"

**Supplementary Figure 1:** Imaging mass cytometry marker visualisation and population clustering.

**A:** UMAPs of representative markers for core microenvironmental populations across all cells. Markers are normalised to the 99.9^th^ percentile of their expression. Diverging scale indicating normalised expression (N.E.). **B:** Schematic of Leiden clustering methods for population derivation. **C:** UMAP of all identified populations generated from Leiden clustering methodology.
