## Supplementary Figure Legend 2 for "High-Dimensional Imaging of Vestibular Schwannoma Reveals Distinctive Immunological Networks Across Histomorphic Niches in *NF2*-related Schwannomatosis"

**Supplementary Figure 2:** Comparison of case-averaged abundance of cell populations across Antoni A and Antoni B regions.

**A:** Comparison of myeloid cell populations. **B:** Comparison of vascular-related populations. **C:** Comparison of T-cell populations**. D:** Comparison of Schwann cell populations. Statistical comparisons were made using unpaired t-tests (for normally distributed data) or Mann Whitney U tests. *=p<0.05.
