## Supplementary Figure Legend 3 for "High-Dimensional Imaging of Vestibular Schwannoma Reveals Distinctive Immunological Networks Across Histomorphic Niches in *NF2*-related Schwannomatosis"

**Supplementary Figure 3:** Comparison of case-averaged abundance of cell populations between genetic disease severity groups.

**A:** Comparison of myeloid cell populations. **B:** Comparison of T-cell populations. **C:** Comparison of Schwann cell populations**. D:** Comparison of vascular-related cell populations. Statistical comparisons were made using unpaired t-tests (for normally distributed data) or Mann Whitney U tests. *=p<0.05.
