## Supplementary Figure Legend 4 for "High-Dimensional Imaging of Vestibular Schwannoma Reveals Distinctive Immunological Networks Across Histomorphic Niches in *NF2*-related Schwannomatosis"

**Supplementary Figure 4:** Generation of cellular neighbourhoods (CNs)

**A:** Elbow plot inferring cluster stability of combined Antoni A and B regions of interest (ROIs). **B:** Dot plot of spatial clusters from combined Antoni A and B ROIs. **C:** Heatmap of spatial cluster distribution across Antoni A and B ROIs, indicating the loss of Antoni B spatial signatures. **D:** Elbow plot inferring cluster stability of Antoni A-only ROIs**. E:** Elbow plot inferring cluster stability of Antoni B-only ROIs.
