## Supplementary Figure Legend 5 for "High-Dimensional Imaging of Vestibular Schwannoma Reveals Distinctive Immunological Networks Across Histomorphic Niches in *NF2*-related Schwannomatosis"

**Supplementary Figure 5:** Effects of bevacizumab-treatment on the microenvironment of vestibular schwannoma.

**A:** Comparison of myeloid cell populations. **B:** Comparison of T-cell populations. **C:** Comparison of Schwann cell populations**. D:** Comparison of vascular-related cell populations. **E:** Elbow plot inferring cluster stability of bevacizumab-inclusive Antoni A-only ROIs. Statistical comparisons were made using unpaired t-tests (for normally distributed data) or Mann Whitney U tests. *=p<0.05.
