## Supplementary Table 1 for "High-Dimensional Imaging of Vestibular Schwannoma Reveals Distinctive Immunological Networks Across Histomorphic Niches in *NF2*-related Schwannomatosis"

**Supplementary Table 1: Antibody clones and sources**

| Acronym | Clone | Source | Identifier |
|---|---|---|---|
| **Recombinant Anti-S100 beta antibody (BSA and Azide free)** | EP1576Y | Abcam | ab215989 |
| **Purified anti-Pan-Cytokeratin Antibody** | AE-1/AE-3 | BioLegend | 914204 |
| **CD45 Monoclonal Antibody** | CD45-2B11 | eBioscience | 14-9457-82 |
| **Recombinant Anti-ICAM1 antibody (BSA and Azide free)** | EP1442Y | Abcam | ab271852 |
| **Recombinant Anti-Granzyme B antibody (BSA and Azide free)** | EPR20129-217 | Abcam | ab219803 |
| **Recombinant Anti-CD16 antibody (BSA and Azide free)** | SP175 | Abcam | ab243925 |
| **Anti-CX3CR1 antibody** | Polyclonal | Abcam | ab8020 |
| **Recombinant Anti-CD11b antibody (BSA and Azide free)** | EP1345Y | Abcam | ab187537 |
| **Recombinant Anti-CD11c antibody (BSA and Azide free)** | EP1347Y | Abcam | ab216655 |
| **CD14 Rabbit mAb** | D7A2T | Cell Signalling Technology | 56082BF |
| **Anti Iba1, Rabbit (for Immunocytochemistry)** | Polyclonal | FUJIFILM Wako Pure Chemical Corp. | 019-19741 |
| **Purified anti-human CD74 Antibody** | LN2 | BioLegend | 326802 |
| **Anti-HLA-DR antibody** | TAL 1B5 | Abcam | ab20181 |
| **CD206/MRC1 Rabbit mAb** | E2L9N | Cell Signalling Technology | 91992 |
| **Purified anti-CD68 Antibody** | KP1 | BioLegend | 916104 |
| **Rat anti-Human CD3** | CD3-12 | Bio-Rad | MCA1477 |
| **Recombinant Anti-CD4 antibody (BSA and Azide free)** | EPR6855 | Abcam | ab181724 |
| **CD8α Monoclonal Antibody** | C8/144B | eBioscience | 14-0085-82 |
| **Purified anti-human CD45RA Antibody** | HI100 | BioLegend | 304102 |
| **Purified anti-human CD45RO Antibody** | UCHL1 | BioLegend | 304202 |
| **Recombinant Anti-FOXP3 antibody (BSA and Azide free)** | 236A/E7 | Abcam | ab96048 |
| **Mouse anti-Human Actin Alpha (Smooth Muscle)** | 1A4 | Bio-Rad | MCA5781GA |
| **CD31/PECAM-1 Antibody - BSA Free** | JC/70A | Novus Biologicals | NB600-562 |
| **Anti-Human Von Willebrand Factor** | Polyclonal | DAKO, Agilent | A0082 |
| **Purified anti-human CD235ab Antibody** | HIR2 | BioLegend | 306602 |
| **Anti-HLA Class 1 ABC antibody** | EMR8-5 | Abcam | ab70328 |
| **Vimentin Antibody - BSA Free** | RV202 | Novus Biologicals | NBP1-97672 |
| **Recombinant Anti-HIF-1 alpha antibody (BSA and Azide free)** | EP1215Y | Abcam | ab210073 |
| **VENTANA PD-L1 Rabbit Monoclonal Primary Antibody** | SP263 | Roche Diagnostics | 07494190001 |
| **Recombinant Anti-Ki67 antibody (BSA and Azide free)** | B56 | Abcam | ab279657 |
| **Phospho-p44/42 MAPK (Erk1/2) (Thr202/Tyr204) XP® Rabbit mAb** | D13.14.4E | Cell Signalling Technology | 4370 |
| **MCT4 Polyclonal antibody** | Polyclonal | Proteintech | #22787-1-AP |
| **Purified anti-mouse/human CD44 Antibody** | IM7 | BioLegend | 103001 |
| **Goat anti-Rabbit IgG (H+L) Cross-Adsorbed Secondary Antibody, Alexa Fluor™ 488** | Polyclonal | ThermoFisher | A-11008 |
